# Integrating AI and causal genetics to prioritize therapeutic targets for aging and age-related diseases

**DOI:** 10.64898/2026.02.13.705676

**Authors:** Geoffrey H. D. Leung, Jianjiu Chen, I. Aylin Ergun, Evgeny Izumchenko, Alex Aliper, Feng Ren, Frank W. Pun, Alex Zhavoronkov

## Abstract

Aging is increasingly viewed as a pathologic process and a principal driver of diverse age-related diseases (ARDs). Framing aging as a disease offers an opportunity to identify therapeutic targets capable of modifying multiple chronic disorders simultaneously. Here, we developed an AI-driven target discovery framework that integrates large-scale multi-omic datasets to prioritise therapeutic targets shared between aging and 12 ARDs across four major disease areas: neurological, inflammatory, metabolic, and fibrotic disorders. We identified 29 high-confidence and 16 previously unrecognized aging-associated targets implicated across selected disease areas, together with convergent pathway perturbations characterized by robust upregulation of interferon and inflammatory signaling, alongside coordinated downregulation of MYC-driven proliferative programs, consistent with heightened inflammatory activation and reduced anabolic activity during aging. Hallmarks of aging assessment revealed chronic inflammation as the most enriched hallmark across aging and ARDs. Mendelian randomization provided genetic causal support for *IL6, IL6R, NLRP3, NOS2, TLR4*, and *GLP1R* in aging-related traits and multiple ARDs, highlighting potential opportunities for drug repurposing. Co-localization analysis further demonstrated a shared genetic signal at the *IL6R* locus between gene expression levels and parental survival. Together, our findings outline a scalable AI-guided multi-omic framework for identifying causal and repurposable therapeutic targets for aging and ARDs.

**Graphical abstract:** 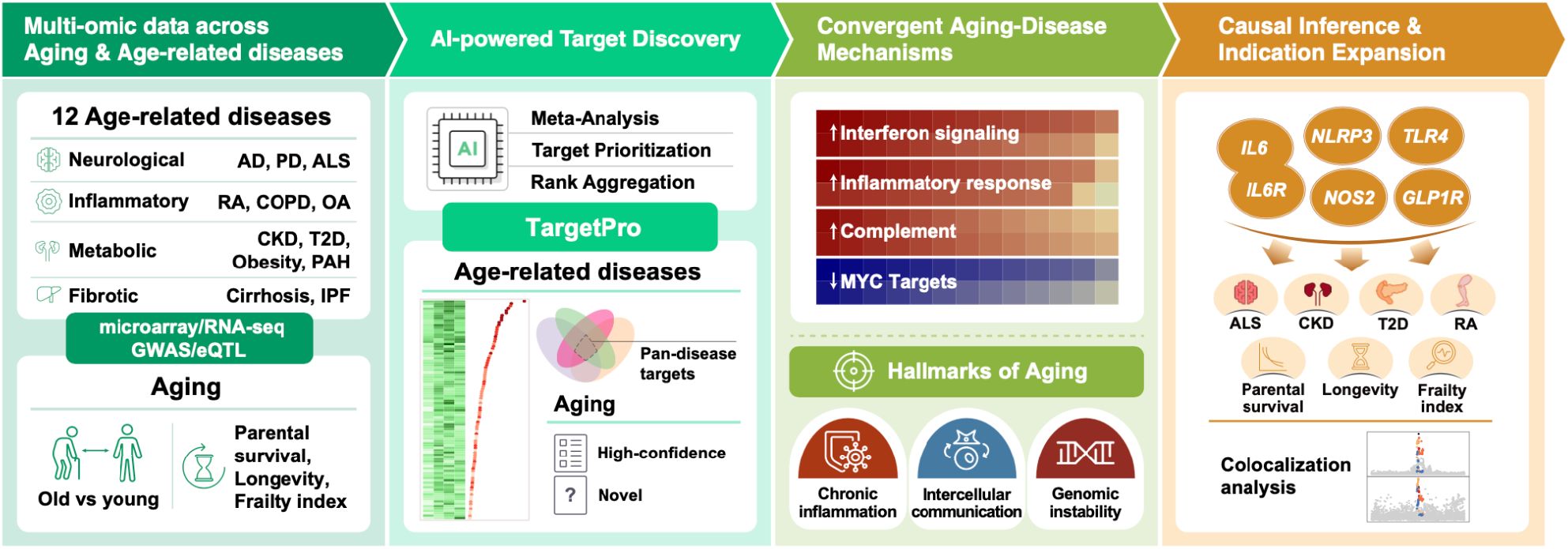

## Introduction

Aging is the dominant risk factor for most chronic diseases, including cardiovascular disorders, cancer, neurodegeneration, and metabolic syndromes. Traditionally viewed as an inevitable biological process, aging is now increasingly understood as a quantifiable and potentially modifiable driver of pathology. Advances in geroscience have demonstrated that the molecular mechanisms underlying aging, such as loss of homeostasis, impaired stress responses, and cellular senescence, are amenable to therapeutic intervention. Considering aging as a disease, or at least as a treatable pathological process, creates the possibility of simultaneously delaying or preventing multiple age-related diseases (ARDs), thereby extending healthspan and reducing the long-term burden on healthcare systems. Reflecting this conceptual shift, aging has been formally recognized in major classification frameworks, including the World Health Organization’s ICD-11 (codes MG2A and XT9T) [1, 2] and the Experimental Factor Ontology (EFO_0022597) [3], underscoring its growing clinical and research relevance.

Recent advances in high-throughput molecular profiling have enabled comprehensive interrogation of aging biology across multiple omic layers, including genomic variation, transcriptomic reprogramming, and epigenetic modifications. These molecular changes interact in complex and non-linear ways to drive both physiological decline and susceptibility to ARDs. However, the scale, heterogeneity, and dimensionality of such datasets present major challenges for conventional analytical approaches, limiting their translational utility. Artificial intelligence (AI), such as deep learning-based methods, offers a powerful solution by enabling the integration of heterogeneous data types, identifying latent patterns across biological systems, and prioritizing therapeutic targets at scale [4-6]. By systematically combining multi-omic data with disease knowledge, AI-driven approaches can uncover shared molecular mechanisms linking aging with diverse pathological outcomes.

In our previous work, we compared transcriptomic signatures of ARDs and non-ARDs as a proxy for aging biology, identifying therapeutic targets enriched across age-related pathologies [7]. In the present study, we reframe aging as a modifiable disease process. By integrating state-of-the-art AI-driven target prioritization methods and causal genetics analyses, we identify circulating therapeutic targets and pathways that mechanistically link aging with multiple ARDs. Our findings highlight both established and novel aging-related targets with geroprotective potential and demonstrate the value of AI-driven multi-omic integration as a scalable framework for systematic target discovery and drug repurposing aimed at extending human healthspan.

## Materials and Methods

### Data collection and processing

Publicly available transcriptomic and DNA methylation datasets were retrieved from the Gene Expression Omnibus (GEO) and ArrayExpress repositories. Only human-derived studies were considered. To ensure sufficient statistical power and analytical robustness, datasets were required to include at least three case samples and three control samples. Analyses were restricted to circulating biospecimens. Raw microarray expression data were normalized by quantile normalization, and RNA-sequencing (RNA-seq) data were normalized by upper-quartile normalization. Genes were filtered to retain protein-coding genes only, as annotated using the biomaRt R package (v2.54.1) [8]. Features with zero coverage in ≥25% of samples were excluded to reduce sparsity and improve downstream statistical stability. In total, 51 datasets spanning aging and 12 age-related diseases were included, comprising 2,839 case samples and 1,886 control samples **(Supplementary Table 1)**.

For Mendelian randomization and colocalization analyses, full GWAS summary statistics for each ARD and two aging-related traits—longevity and frailty index—were obtained from the GWAS Catalog [9] **(Supplementary Table 2)**. In the longevity GWAS, cases were defined as individuals surviving to or beyond the 90th percentile of age within the cohort, while controls were defined as individuals who died before the 60th percentile of age [10]. Frailty index was defined as the number of health deficits during an individual’s life-course such as immobility, disability, and hospitalization [11]. In addition, parental survival was included as a proxy for lifespan, under the assumption of shared genetic effect sizes between maternal and paternal survival [12]. To minimize population stratification, only GWAS conducted in European-ancestry cohorts were included. For traits with multiple eligible GWAS, the study with the largest sample size and most comprehensive variant coverage was retained. Expression quantitative trait loci (eQTL) data were obtained from the eQTLGen Phase I dataset [13]. To ensure consistency across genetic analyses, all GWAS and eQTL genomic coordinates were harmonized to the GRCh37 reference genome using the UCSC’s LiftOver tool where necessary [14].

### Quality control

Quality control (QC) was applied to each dataset prior to downstream analyses. Initial assessment included visualization of sample distributions using boxplots and principal component analysis to identify batch or plate effects and detect potential outlier samples arising from technical artefacts.

Datasets exhibiting clear technical outliers or non-biological clustering patterns were examined further prior to inclusion. For ARD datasets, additional steps were taken to account for potential confounding by chronological age, as age-associated molecular changes in gene expression and DNA methylation could bias disease–control comparisons. The contribution of age to feature-level variability was quantified using the variancePartition R package (v1.28.9) [15], which estimates the proportion of variance explained by each covariate. Age was included as a covariate in linear models if it explained more than 5% of the variance in over 25% of features within a dataset, and adjusted residuals were derived using the lm() function in R. The resulting age-adjusted feature matrix was used for all subsequent analyses. The same QC framework was applied to other available covariates across all datasets, including both aging and age-related disease cohorts. Potential confounders such as sex, geographic region, and ethnicity were evaluated for their contribution to feature-level variance. Where these covariates exceeded the predefined variance thresholds, their effects were similarly regressed out prior to downstream analyses. All QC-filtered and covariate-adjusted matrices were subsequently uploaded to PandaOmics for subsequent AI-driven target discovery by TargetPro.

### Case-control comparisons and target prioritization

Case-control comparison was performed separately for each dataset in each condition using *limma* implemented in PandaOmics. For aging datasets, cases were defined as individuals aged ≥60 and controls as those aged ≤50. For 1 aging dataset (GSE16717), long-lived individuals (mean age = 93.4 years) were treated as cases where non-long-lived individuals (mean age = 61.9 years) from the same families were included as controls. For each condition, results from all eligible datasets were subsequently combined using meta-analysis to derive consensus disease-associated effect estimates across studies. P values were adjusted for multiple testing using the Benjamini–Hochberg (BH) procedure, and genes with BH-adjusted P values (P_BH_) < 0.05 (i.e., leading to an FDR < 0.05) were considered significantly dysregulated. Target prioritization was performed using the AI-driven target identification tool TargetPro available on PandaOmics [16, 17]. Briefly, TargetPro integrates evidence from 22 independent models, which are subsequently combined using an XGBoost ensemble to rank genes according to their likelihood of causal involvement in a given disease. These models leverage multiple evidence types, including clinical annotations and prior knowledge of target–disease associations, such as whether a gene has been investigated in clinical trials or approved therapies for the disease of interest. Since aging has only recently been classified as a disease and lacks standardized clinical trial endpoints, direct clinical evidence for aging-targeted interventions remains sparse. To address this limitation, we incorporated a curated list of aging-associated genes from the GenAge database (302 genes) [18] as a surrogate source of prior biological evidence for high-confidence aging-related targets. This gene set was used to inform and calibrate target prioritization for aging within the TargetPro framework, enabling systematic ranking of candidate targets despite the limited availability of clinical aging-specific data. In contrast, novel aging-related targets were defined as those not annotated in the GenAge database.

### Pathway enrichment analysis

Gene Set Enrichment Analysis (GSEA) was performed on aging and each ARD using ranked gene lists derived from the meta-analyzed effect estimates (β), representing overall transcriptomic changes between cases and controls. Fifty human hallmark gene sets were obtained from the Molecular Signatures Database (MSigDB) via the msigdbr R package (v25.1.1) [19]. GSEA was conducted using the fgsea R package (v1.24.0) [20]. Gene sets with positive normalized enrichment scores (NES > 0) were interpreted as transcriptionally activated, whereas those with negative scores (NES < 0) were considered inhibited. In addition to GSEA, over-representation pathway enrichment analysis was performed to identify biological pathways enriched among gene sets commonly implicated across disease areas, as well as among high-confidence and novel aging-specific target genes. Reactome-annotated pathways were analyzed using the R package ReactomePA (v1.42.0) [21]. For both hallmark gene sets and Reactome pathways, multiple testing correction was applied using the Benjamini–Hochberg procedure, and pathways with P_BH_ < 0.05 were considered significantly enriched.

### Robust rank aggregation

Robust rank aggregation (RRA) was applied to identify hallmark gene sets and target genes that were consistently prioritized higher than expected across multiple ARDs [22]. RRA was performed using the *RobustRankAggregation* R package (v1.2.1) [22]. Briefly, ranked lists of hallmark gene sets (50 gene sets) or target genes (19,291 genes) were generated for each ARD. RRA was conducted in a two-stage manner. First, ranked lists were combined within each disease area to identify gene sets and targets that were consistently enriched or prioritized across area-specific ARDs. Subsequently, ranked lists from all ARDs were aggregated to identify pan-ARD hallmark gene sets and target genes shared across disease areas. RRA assigns a score to each gene set or target gene that reflects the probability of observing its aggregated rank under a null model of random ordering, with lower scores indicating stronger and more consistent prioritization. For visualization and interpretability, RRA scores were –log_10_-transformed, such that higher aggregated scores correspond to stronger evidence of consistent enrichment or prioritization across ARDs.

### Hallmarks of aging assessment

Assessment on the associated hallmark(s) of aging was performed for each target gene based on their annotated gene ontology (GO) terms retrieved from biomaRt and literature by mapping the keywords as previously described [23]. A gene was defined as associated with a particular hallmark if it had ≥1 GO term matched to the keyword(s) of that corresponding hallmark.

### Mendelian randomization

Cis-acting eQTLs (cis-eQTLs) were used as instrumental variables in Mendelian randomization (MR) analyses to minimize horizontal pleiotropy and enhance biological interpretability. Cis-eQTL variants were first filtered using a genome-wide significance threshold (P < 5×10^−8^). Linkage disequilibrium (LD) clumping was then performed to retain approximately independent variants, using PLINK (v1.9) [24] with the European reference panel from the 1000 Genomes Project (phase 3). Variants were clumped using an LD threshold of r^2^ < 0.1 within a ±100 kb window. The resulting LD-pruned cis-eQTL variants were used as instruments in two-sample MR analyses using the R package TwoSampleMR (v0.5.7) [25] to test for putative causal effects of gene expression on the risk of each ARD and aging-related trait. Summary statistics were obtained from independent eQTL and GWAS datasets. For genes with multiple instrumental variants, causal estimates were primarily derived using the inverse-variance weighted (IVW) method. When only a single variant remained after significance filtering and LD clumping, causal effects were estimated using Wald’s ratio. MR associations with nominal P < 0.05 were considered statistically significant. Colocalization analysis was conducted for each gene–trait pair showing significant MR evidence, using the approximate Bayes factor method implemented in the *coloc* R package (v5.2.3) [26]. Posterior probabilities were computed to assess whether the eQTL and disease association signals were consistent with a shared causal variant. Strong colocalization was defined as PP.H4 > 0.8, and moderate colocalization as PP.H4 > 0.6. Prior probabilities were set to p_1_ = 1×10^−4^, p_2_ = 1×10^−4^, and p_12_ =1×10^−5^.

## Results

### Targets and pathways implicated in age-related diseases

A total of 12 age-related diseases (ARDs) from 4 disease areas were included in this study, including neurological diseases: Alzheimer’s disease (AD), Amyotrophic lateral sclerosis (ALS), Parkinson’s disease (PD); inflammatory diseases: chronic obstructive pulmonary disease (COPD), Osteoarthritis (OA), Rheumatoid arthritis (RA); metabolic diseases: Chronic kidney disease (CKD), Obesity, Pulmonary arterial hypertension (PAH), Type II diabetes (T2D); and fibrotic diseases: Cirrhosis of liver (Cirrhosis), Idiopathic pulmonary fibrosis (IPF) (**Supplementary Table 1**). Gene Set Enrichment Analysis (GSEA) was performed on each ARD using the meta-analyzed effect estimates obtained from differential expression analysis to identify perturbed gene sets. Out of the 50 hallmark gene sets tested, IPF showed the most significant dysregulation of 25 (50%) gene sets (FDR < 0.05), followed by PAH (20 gene sets, 40%), and obesity (18 gene sets, 36%) (**Supplementary Figure 1**). Robust rank aggregation (RRA) identified hallmark gene sets that were consistently perturbed across diseases within each disease area (**Figure 1A**). This revealed a core set of consistently activated pathways, including interferon gamma response, inflammatory response, complement, and apoptosis. In contrast, MYC targets v1 was broadly downregulated across most ARDs, suggesting suppression of proliferative and biosynthetic programs in chronic disease contexts (**Figure 1A**).

**Figure 1.**
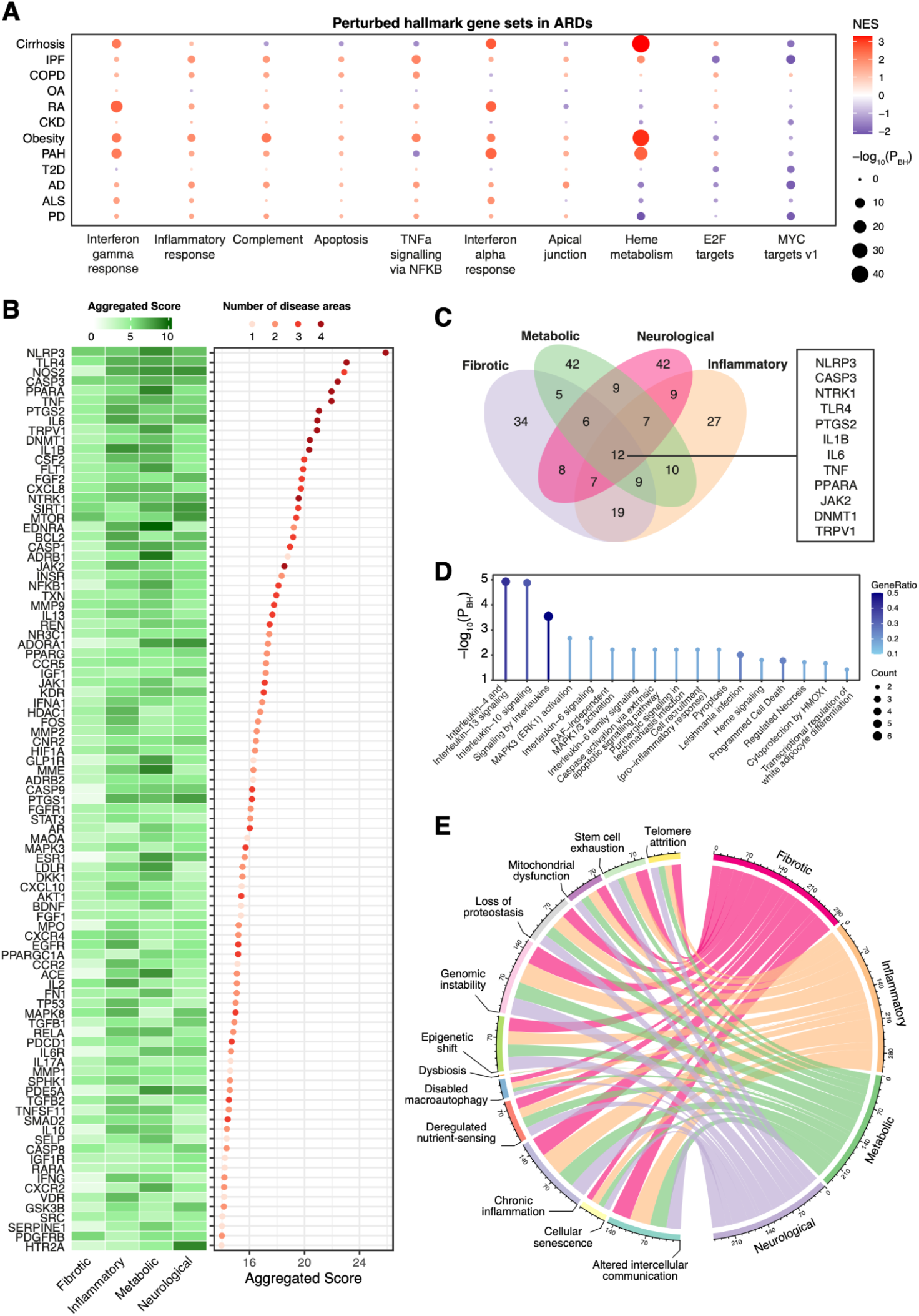
Transcriptional changes and targets implicated in age-related diseases. **A.** The 10 most significantly perturbed hallmark gene sets ranked by robust rank aggregation (RRA) across ARDs. Red, NES > 0; Blue, NES < 0. **B**. Ninety-three target genes that were highly ranked across ARDs by RRA. The heatmap shows their aggregated score in each disease area. The dotplot shows the overall aggregated score for each gene across all ARDs and the number of disease areas involved. **C**. Overlapping the number of prioritized targets between disease areas. Twelve targets highly ranked across all areas were listed in the box. **D**. Pathway enrichment analysis of the 12 target genes highly ranked across disease areas shown in **C. E**. Associations between hallmarks of aging and disease areas based on the targets prioritized. NES, Normalized enrichment score; P_BH_, Benjamini-Hochberg-corrected P value.

Target prioritization was performed for each disease to identify the most promising therapeutic gene candidates based on 22 machine learning models (see **Materials and Methods**). Ranked gene lists were aggregated using RRA within each disease area (top 100 genes per area; union = 246 genes) and across all ARDs, yielding 93 target genes that were prioritized higher than expected across diseases (**Figure 1B**). Among these, *NLRP3* and *TLR4* emerged as the most consistently prioritized targets, which were implicated across all disease areas, followed by *NOS2*, which was highly ranked in inflammatory, metabolic, and neurological diseases. Twelve target genes were commonly enriched across all disease areas, including *NLRP3, CASP3, NTRK1, TLR4, PTGS2, IL1B, IL6, TNF, PPARA, JAK2, DNMT1*, and *TRPV1* (**Figure 1C)**. Pathway enrichment analysis of these 12 pan-disease genes revealed 17 significantly enriched pathways (FDR < 0.05). Interleukin-4 and Interleukin-13 signaling was the most enriched pathway (P_BH_ = 1.2×10^−5^), followed by Interleukin-10 signaling (P_BH_ = 1.3×10^−5^). Additional enriched pathways included MAPK signaling, programmed cell death, immune regulation, and adipocyte differentiation, highlighting shared inflammatory and metabolic mechanisms across ARDs (**Figure 1D**).

We next assessed the relationship between prioritized targets and aging via their associations with hallmarks of aging (see **Materials and Methods**). Of the 93 genes tested, 91 (98%) were associated with at least one hallmark. As expected, inflammatory diseases showed the strongest enrichment for the hallmark of chronic inflammation, with 52 of 63 inflammatory-associated targets mapping to this process (P = 1.5×10^−3^, hypergeometric test). Across all disease areas, chronic inflammation was the most frequently implicated hallmark (67 genes, 72%), followed by altered intercellular communication (65 genes, 70%) and genomic instability (64 genes, 69%) (**Figure 1E**). Together, these findings demonstrate important involvement of aging-related biological processes across ARDs, with inflammation emerging as a central shared axis.

### Targets and pathways implicated in aging

To characterize transcriptomic changes associated with aging, four independent datasets were analyzed comparing older and younger individuals (see **Materials and Methods**). Meta-analysis revealed 878 genes that were significantly differentially expressed in older adults, with 312 and 566 genes upregulated and downregulated, respectively (FDR < 0.05, **Figure 2A**). The most significantly upregulated genes included *FLVCR2, SORT1, ANXA2, LGALS1*, and *FCGR1A*, while *LRRN3, CD248, CR2, NOG*, and *SLC4A10* were among the most significantly downregulated. GSEA revealed 33 significantly enriched hallmark gene sets in aging. Interferon gamma response (P_BH_ = 3.9×10^−32^) and interferon alpha response (P_BH_ = 8.4×10^−25^) were the most strongly enriched pathways (**Figure 2B**), indicating pronounced activation of immune and inflammatory signaling with age. Notably, these pathways mirrored those consistently dysregulated across ARDs (**Figure 1A**), highlighting shared transcriptional programs between aging and disease states.

**Figure 2.**
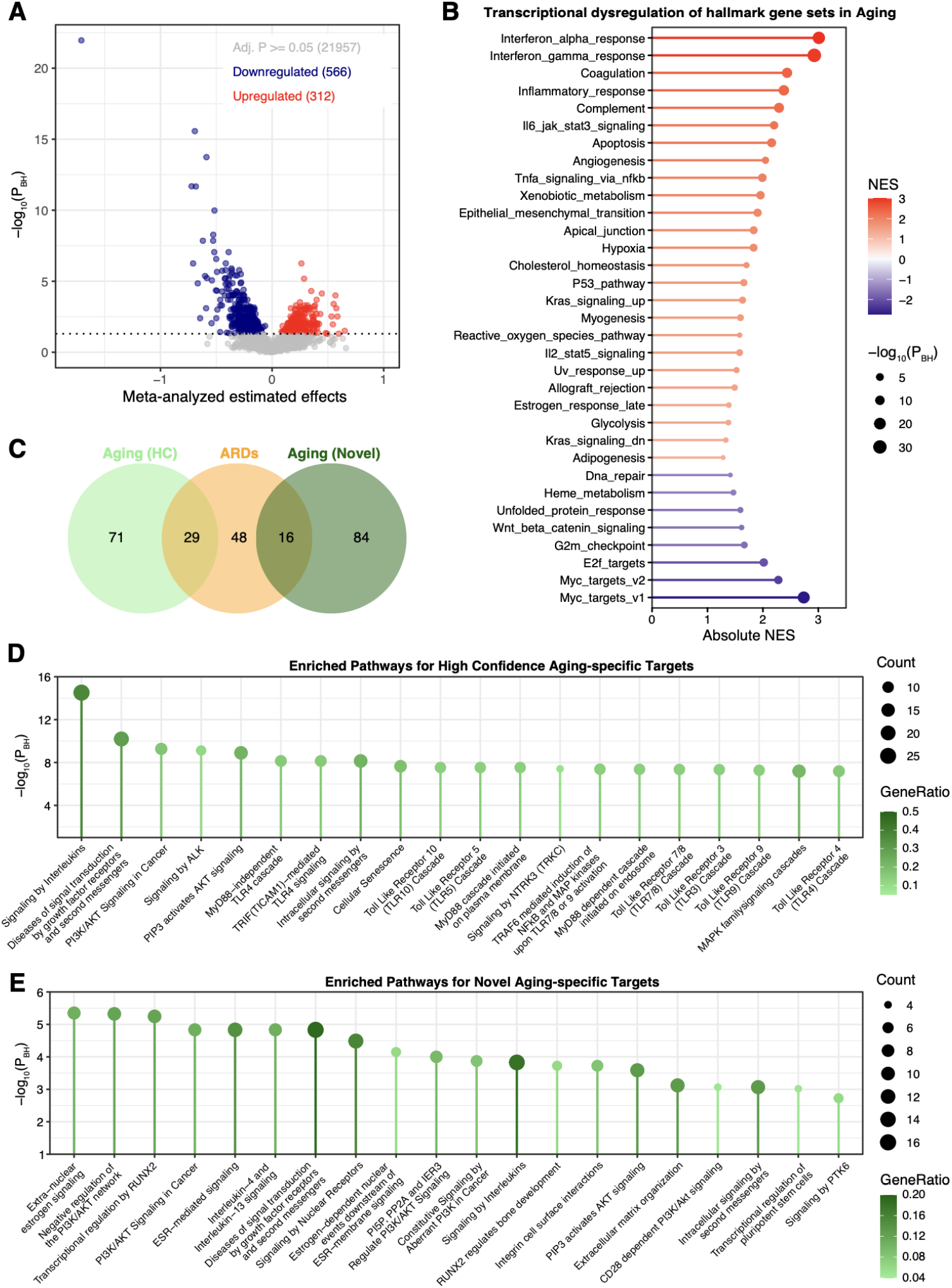
Transcriptional changes and targets implicated in aging. **A.** Volcano plot showing the meta-analyzed estimated effects (log_2_ fold change) across the 4 transcriptomic datasets of aging. Each point represents one gene. Red, significantly upregulated; Blue, significantly downregulated; Grey; not significantly dysregulated at P_BH_>0.05. **B**. Dysregulation patterns of 33 significantly perturbed hallmark gene sets in aging. **C**. Venn diagram showing the number of overlapped targets between aging and age-related diseases, and aging-specific target genes. **D**. Pathways enriched for high-confidence aging-specific target genes. **E**. Pathways enriched for novel aging-specific target genes. HC, High confidence; NES, Normalized enrichment score; P_BH_, Benjamini-Hochberg-corrected P value; ARDs, Age-related diseases.

Target prioritization identified a set of genes previously implicated in aging, as included in the established database of human aging genes (GenAge). These included *CDKN2A, PTPN11, JAK2, SOD1, VEGFA* (**Supplementary Table 3**), which we define as high-confidence aging targets. To increase discovery potential, we additionally prioritized genes not present in the GenAge database, generating a list of novel aging targets. Highly ranked candidates included *CXCL8, CD19, IL3, CASP3*, and *TLR4* (**Supplementary Table 4**). Aging-specific targets were defined as those prioritized in aging but not present in the pan-ARD target list (**Figure 1B**), yielding 71 high-confidence and 84 novel aging-specific targets (**Figure 2C**).

Pathway enrichment analysis on the high-confidence aging-specific targets identified signaling by interleukins as the most enriched pathway (P_BH_ = 3.0×10^−15^), alongside enrichment of PI3K/AKT signaling, cellular senescence, NTRK signaling, growth factor signaling, and multiple Toll-like receptor pathways (**Figure 2D, Supplementary Table 5**). For novel aging-specific targets, extra-nuclear estrogen signaling was the most enriched pathway (P_BH_ = 4.5×10^−6^), with additional enrichment in PI3K/AKT, interleukin signaling, RUNX2 transcriptional programs, extracellular matrix organization, pluripotency-related transcription, and PTK6 signaling (**Figure 2E, Supplementary Table 6**). These results highlight aging-specific regulatory programs extending beyond classical inflammatory pathways to include hormonal, developmental, and stem cell– associated signaling.

### Commonly perturbed pathways and targets implicated in aging and age-related diseases

To directly compare aging and ARDs at the pathway level, we examined enrichment patterns of the 33 aging-associated hallmark gene sets across ARDs. Most gene sets exhibited consistent dysregulation across diseases, including activation of complement, TNF-α signaling, inflammatory response, and interferon pathways (**Figure 3A**). In contrast, DNA repair was downregulated in aging but showed weak, non-significant upregulation across most ARDs, suggesting potential divergence between physiological aging and disease-associated compensatory mechanisms. Correlation analysis of normalized enrichment scores demonstrated strong concordance between aging and neurological diseases (PD: ρ = 0.93; AD: ρ = 0.86; ALS: ρ = 0.73), indicating shared transcriptional architectures (**Figure 3B**). Moderate-to-strong correlations were also observed for several other diseases, whereas cirrhosis (ρ = 0.15) and OA (ρ = −0.03) were poorly correlated with aging, reflecting possibly limited statistical power, tissue heterogeneity, or disease-specific biology.

**Figure 3.**
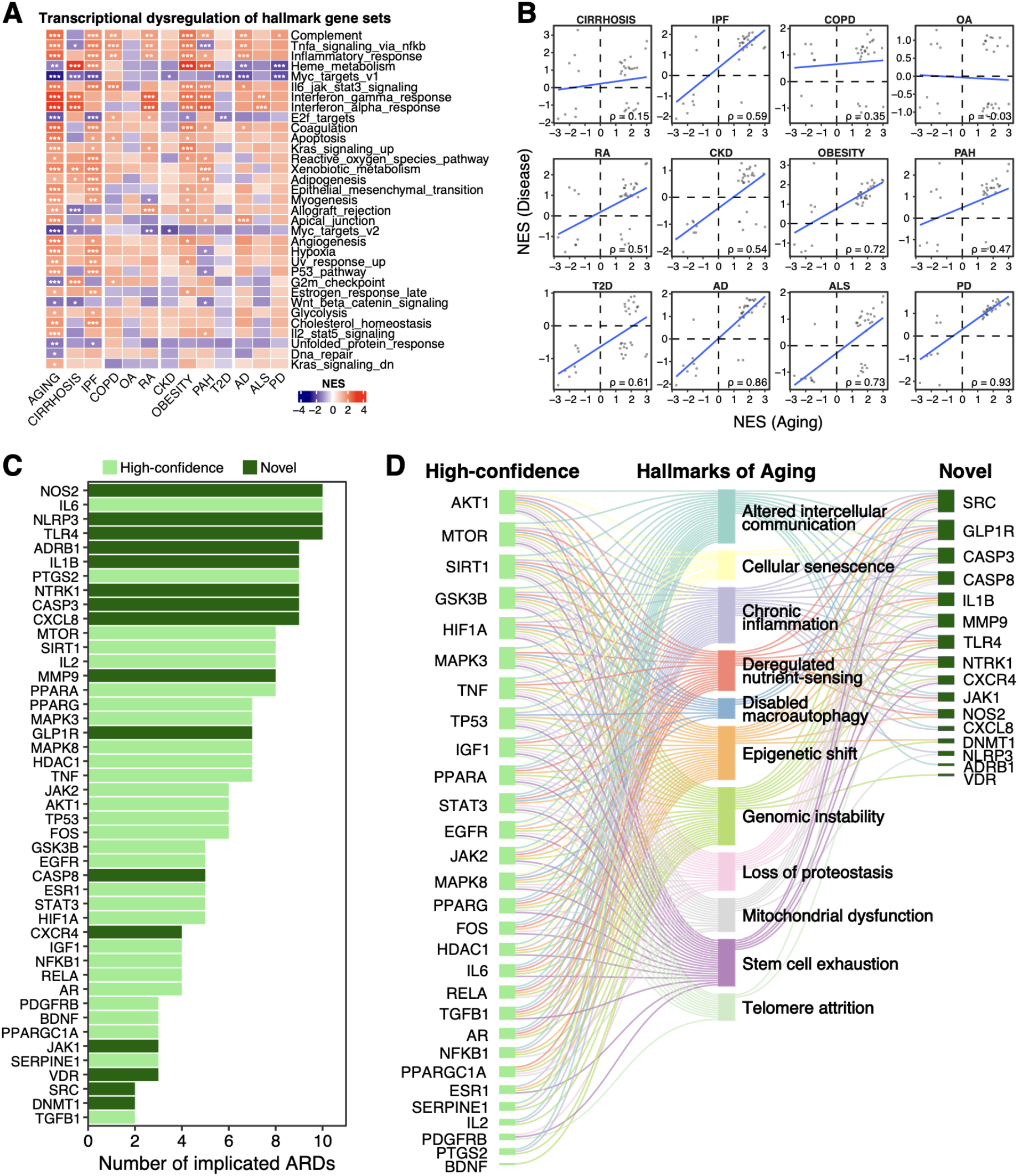
Shared pathways and targets between aging and age-related diseases. **A.** Transcriptional dysregulation patterns of 33 hallmark gene sets in aging and ARDs that were significantly perturbed in aging. *P_BH_<0.05, **P_BH_<0.01, ***P_BH_<0.001. **B**. Correlation of the normalized enrichment scores for all hallmark gene sets between aging and ARDs. Each point represents one gene set. Blue line indicates the best-fit line obtained from linear regression. **C**. Forty-five commonly implicated target genes in aging and ARDs. **D**. Sankey plot showing the associations between 45 commonly implicated targets (29 high-confidence and 16 novel aging targets) and hallmarks of aging. NES, Normalized enrichment score; P_BH_, Benjamini-Hochberg-corrected P value.

Forty-five targets (29 high-confidence and 16 novel aging-related targets) were prioritized in aging and ARDs (**Figure 2C**). Among these, *NOS2, IL6, NLRP3*, and *TLR4* were implicated in 10 ARDs (**Figure 3C**). All of these targets were associated with at least 1 hallmark of aging (**Figure 3D**). Among the high-confidence targets, *AKT1, MTOR*, and *SIRT1* were associated with 11 hallmarks, while *SRC* and *GLP1R*, among the novel targets, were associated with 10 and 9 hallmarks, respectively. Chronic inflammation was again the most enriched hallmark (linked to 38 genes, 84%), reinforcing its central role in both aging and disease.

### Mendelian randomization identifies genes causally linked to aging-related traits

To assess potential causal relationships between the prioritized targets through our discovery pipeline and aging, we performed Mendelian randomization (MR) analyses using cis-eQTL instruments (see **Materials and Methods**). Aging-related traits included parental survival [12], longevity [10], and frailty index [11] (see **Materials and Methods**). We selected 1 high-confidence (*IL6*) and 3 novel aging genes (*NOS2, NLRP3* and *TLR4*), which collectively showed prioritization in 10 ARDs. We additionally included *GLP1R* given primarily its strong enrichment across 9 hallmarks of aging in the novel target list and its prioritization in 7 ARDs. Because IL-6 exerts its biological effects through IL-6 receptor (IL6R), the pharmacological target of approved IL-6 pathway inhibitors, we also evaluated *IL6R* as a mechanistically linked candidate.

*IL6R, NOS2, NLRP3* and *TLR4* showed significant MR associations with at least 1 aging-related trait. Increased *IL6R* expression was associated with poorer parental survival (β = −0.10, P = 1.6×10^−4^, **Figure 4A**), showing its association with aging in addition to prior evidence linking IL-6 signaling to inflammaging [27-29]. *NOS2* expression was positively associated with frailty index (β = 0.048, P = 0.018, **Figure 4B**). *NLRP3* was positively associated with longevity (β = 0.077, P = 0.014) and negatively associated with frailty index (β = −0.018, P = 6.6×10^−3^). *TLR4* showed modest association with parental survival (β = 0.008, P = 0.048) but a stronger negative association with longevity (β = −0.066, P = 1.1×10^−4^). Colocalization analysis revealed strong evidence for a shared causal variant between *IL6R* expression and parental survival (PP.H4 = 0.84; **Figure 4C**), driven by rs10908839 (SNP.PP.H4 = 0.99), supporting a shared genetic architecture underlying IL-6 signaling and survival.

**Figure 4.**
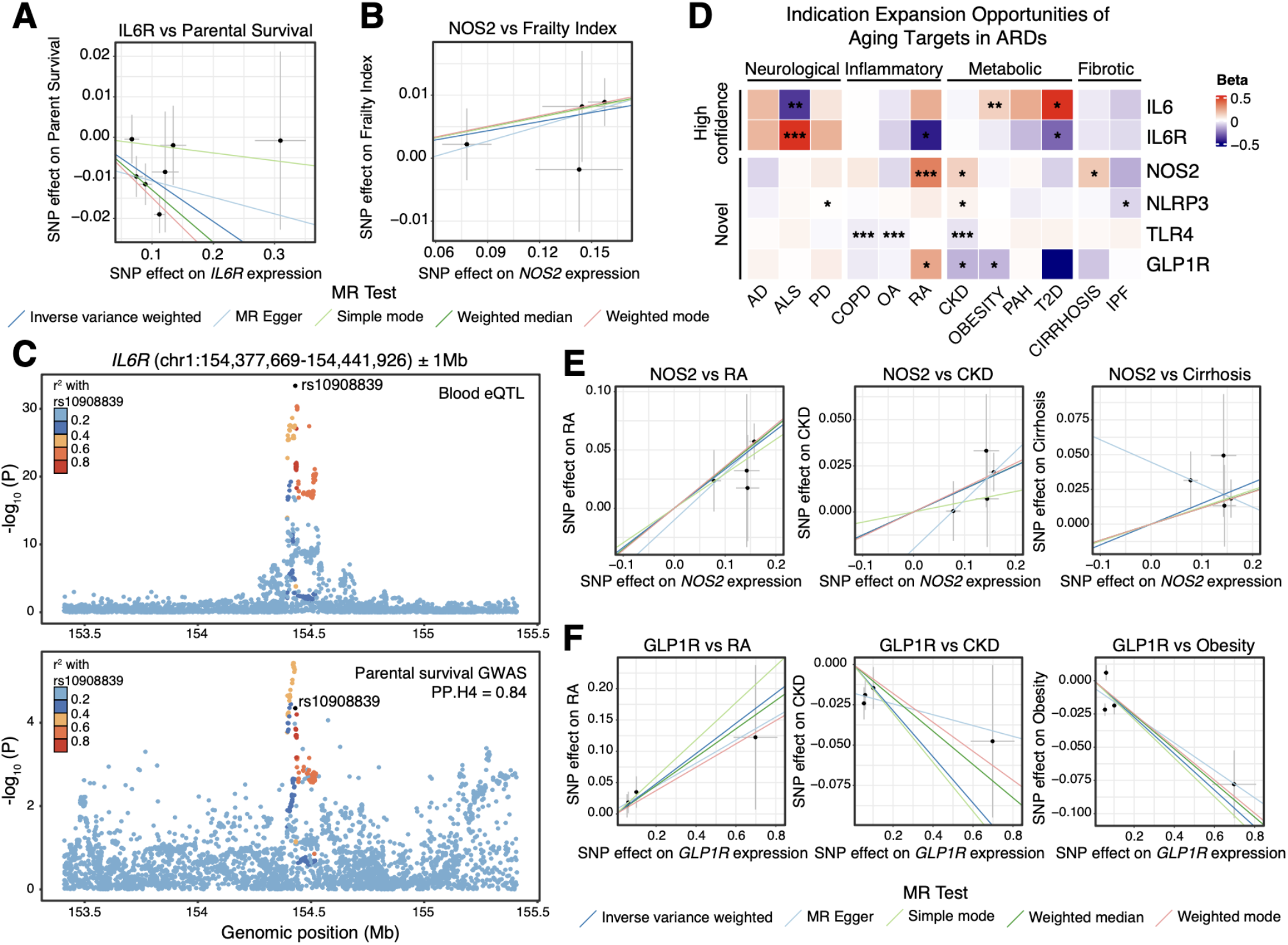
Genetic associations of prioritized dual-purpose targets in age-related diseases. **A.** Mendelian randomization (MR) suggests a significant causal relationship between the genetic variants of *IL6R* and parental survival. **B**. Significant MR results showing the causal relationship between *NOS2* expression and frailty index. **C**. Colocalization analysis showing strongly shared genetic signals between *IL6R* expression and parental survival at the *IL6R* locus. **D**. Indication expansion opportunities of prioritized aging targets in ARDs; *P_BH_<0.05, **P_BH_<0.01, ***P_BH_<0.001. **E**. Significant causal relationships between *NOS2* and RA, CKD, and cirrhosis of liver suggested by MR. **F**. Significant causal relationships between *GLP1R* and RA, CKD, and Obesity.

### Indication expansion opportunities across ARDs

We next evaluated indication expansion opportunities by testing causal associations between shortlisted targets and ARDs. All nominated targets showed significant MR associations with at least one ARD (**Figure 4D**). Specifically, *IL6* (β = –0.39, P = 9.9×10^−3^) and *IL6R* (β = 0.55, P = 3.6×10^−4^) were significantly associated with ALS risk, while *NOS2* showed associations with RA, CKD, and cirrhosis (**Figure 4E**), suggesting therapeutic potential for NOS2 inhibition. *NLRP3* showed suggestive associations with CKD and IPF, and *TLR4* demonstrated strong causal associations with CKD, COPD, and OA. Finally, *GLP1R* was associated with RA, obesity, and CKD (**Figure 4F**), consistent with established metabolic effects of GLP-1R agonists, while highlighting disease-specific complexities. Notably, all four novel targets showed significant MR associations with CKD, suggesting convergent inflammatory and metabolic mechanisms in renal aging and disease. Colocalization analysis further revealed moderately shared signals between gene expression levels and disease risk of RA at the *NOS2* locus (PP.H4 = 62.1%, **Supplementary Figure 2**), supporting shared genetic mechanisms underlying *NOS2* regulation and RA susceptibility.

## Discussion

Aging is a major risk factor for a wide spectrum of chronic conditions, yet the extent to which shared molecular processes underlie both intrinsic ageing biology and clinically defined age-related diseases (ARDs) remains incompletely understood. By integrating transcriptomic meta-analysis, AI-driven target prioritization, and genetic-based causal inference, we provide a systematic framework to identify shared and aging-specific mechanisms with translational relevance.

Across the 12 ARDs examined, we observed widespread activation of hallmark pathways related to interferon signaling, complement activation, inflammatory response, and apoptosis. Importantly, these signatures were not confined to traditionally immune-driven diseases (e.g., RA) but extended to neurodegenerative (AD, PD, ALS), metabolic (obesity, T2D, CKD, PAH), and fibrotic diseases (IPF, cirrhosis), indicating shared systemic immune perturbation across clinically distinct phenotypes. This aligns with the geroscience concept of “inflammaging,” describing chronic, low-grade inflammatory activation as a key feature of aging that contributes to multimorbidity and mortality risk [30]. Our pan-disease meta-signature further emphasized consistent perturbation of interferon-γ response and complement activation across disease areas, reinforcing immune dysregulation as a common denominator across ARDs. The broad downregulation of MYC targets across diseases is also notable given MYC’s roles in proliferation, metabolism, and immune cell function; suppression of MYC-regulated programs may reflect chronic stress adaptation, altered immune cell composition, or disease-associated metabolic reprogramming shared across ARDs.

Deep learning-based target prioritization followed by robust rank aggregation yielded a relatively compact set of 93 pan-ARD targets, led by innate immune sensors and inflammatory mediators including *NLRP3* and *TLR4*, and supported by canonical cytokines (*IL1B, IL6, TNF*). The convergence of these targets across disease areas suggests that upstream inflammatory sensing and downstream cytokine signaling are not merely correlates of individual diseases but may represent shared, potentially targetable processes across multiple ARDs. This observation is consistent with a growing literature implicating NLRP3 inflammasome activation and innate immune signaling in aging-associated chronic inflammation and degenerative pathology [31]. Beyond classical inflammatory mediators, the enrichment of metabolic and regulatory targets such as *PPARA, DNMT1*, and *JAK2* supports a view of ARDs as diseases of integrated immune-metabolic dysfunction, in which inflammatory activation, epigenetic regulation, and cellular stress responses converge. Pathway enrichment of the 12 consensus targets highlighted interleukin signaling pathways (including IL-4/IL-13 and IL-10 signaling), MAPK activation, apoptosis, and adipocyte differentiation, pointing to a cytokine-driven backbone that links immune regulation with tissue remodeling and metabolic dysregulation. When mapped to the hallmarks of aging, the majority of pan-ARD targets aligned with chronic inflammation, altered intercellular communication, and genomic instability, supporting the concept that many ARDs reflect accelerated or dysregulated manifestations of aging biology rather than independent disease entities.

Our aging meta-analysis revealed broad transcriptional changes, with interferon-γ and interferon-α responses being the most significantly enriched hallmark pathways. The prominence of interferon programs in aging is consistent with emerging evidence that age-associated immune remodeling includes heightened interferon-related transcriptional activity in human blood and immune cells, and that such programs can be observed across inflammatory contexts [32-34]. Notably, these interferon signatures mirrored the most conserved pathways observed across ARDs in our analysis, strengthening the hypothesis that immune activation is not purely secondary to disease, but may represent an aging-associated baseline state that predisposes tissues to maladaptive responses. In addition to rediscovering established aging regulators (e.g., CDKN2A, JAK2, VEGFA), our stratification into high-confidence and novel aging targets suggested that aging-associated remodeling encompasses both well-characterized senescence-linked pathways and underexplored immune and signaling signatures. Enrichment of PI3K/AKT, growth factor signaling, Toll-like receptor pathways, extracellular matrix programs, and estrogen-linked signaling underscores the complex nature of aging biology and points to therapeutic opportunities beyond direct cytokine blockade. Direct comparison of pathway signatures revealed strong concordance between aging and neurological diseases (particularly PD and AD), supporting that neurodegenerative disease may represent an intensified or context-specific deployment of aging-associated immune and stress programs. This is in line with the steep age-dependence of neurodegenerative conditions and the expanding literature linking neuroinflammation and innate immunity to neurodegeneration [35]. In contrast, osteoarthritis and cirrhosis showed weak correlations with aging signatures in our dataset. This divergence may reflect limited statistical power (notably for OA where only 1 dataset was included), heterogeneity of circulating signatures relative to tissue-local pathology, or strong contributions of non-aging-dominant drivers (e.g., biomechanical loading in OA; toxin-, metabolic-, or infection-driven injury in cirrhosis) [36-38]. Future work incorporating larger cohorts, tissue-resolved profiling, and longitudinal sampling will be essential to distinguish true biological divergence from these constraints.

The integration of human genetics in our framework strengthened causal inference. Using cis-eQTL instruments, MR and colocalization analyses provided orthogonal support for several prioritized targets and clarified which candidates may reflect causal regulation rather than reactive transcriptional changes. The expression of *IL6R*, the receptor for our top hit, *IL6*, showed a significant association with parental survival alongside strong colocalization, consistent with prior genetic evidence linking IL-6 signaling to cardiovascular, immune-related outcomes and longevity-related phenotypes [39-41]. These findings support the argument that IL-6 pathway modulation is relevant not only for inflammatory disease endpoints but also for aging-related functional decline. However, parental survival is an indirect proxy for lifespan and may be influenced by shared environmental, socioeconomic, and behavioral factors, which could partially confound genetic interpretation. Consistently, Rosa et al. previously reported that genetically proxied higher soluble IL6R, interpreted as reduced IL-6 signaling, was associated with improved parental survival [42]. Despite this, experimental studies of IL6R inhibition have not demonstrated robust lifespan extension despite reducing inflammation [43], possibly because IL-6 signaling influences disease susceptibility and functional decline more strongly than organismal lifespan, or because compensatory inflammatory pathways limit survival benefits. Nevertheless, our results support a context-dependent role for IL6R in aging, where modulation of IL-6 signaling may improve healthspan and disease outcomes without necessarily extending lifespan.

MR evidence also supported associations of *NOS2, NLRP3*, and *TLR4* with aging-related traits and/or ARDs, although effect directionality and trait specificity varied, underscoring the context-dependent roles of innate immune signaling and the need for careful translational interpretation. Our genetic analyses further highlighted indication expansion opportunities, with convergence of causal signals in chronic kidney disease across several targets, suggesting the kidney as a nexus where inflammatory, metabolic, and vascular aging processes intersect. For NOS2, moderate colocalization with RA provides genetic support for shared regulatory mechanisms linking nitric oxide signaling and inflammatory disease susceptibility. Historical pharmacological development illustrates the complexity of translation, as selective iNOS inhibition (e.g., GW-274150) showed limited efficacy in migraine and asthma [44, 45]. Nonetheless, broader targeting of nitric oxide pathways has been proposed across cardiovascular disease [46], cancer [47], erectile dysfunction [48], and neurodegeneration [49], supporting continued interest in NOS2-related mechanisms across ARDs and potentially aging.

Our study has several limitations. First, our transcriptomic analyses were restricted to circulating biospecimens to maximize cross-study comparability, but this design may underrepresent tissue-local mechanisms that are critical in certain ARDs (e.g., synovium in OA/RA, liver in cirrhosis, lung parenchyma in IPF). Second, heterogeneity across public datasets (platform differences, batch effects, clinical covariates, and case definitions) is unavoidable; while we applied QC procedures and meta-analysis to mitigate these issues, residual heterogeneity may influence specific disease-level signals. Third, AI-driven target prioritization depends partly on pre-existing knowledge (e.g., clinical annotations), which can bias rankings toward well-studied targets; our separation into high-confidence versus novel aging targets partly addresses this, but does not fully eliminate ascertainment bias. Fourth, MR and colocalization analyses have inherent assumptions and constraints: cis-eQTL instruments reflect gene regulation in specific tissues and contexts (blood in this study), may not capture disease-relevant cell states, and can still be influenced by LD structure or pleiotropy; additionally, our analyses were restricted primarily to European-ancestry GWAS, limiting generalizability. Finally, nominal MR thresholds followed by colocalization provide a pragmatic discovery-validation workflow, but future work should incorporate more extensive sensitivity analyses, triangulation with pQTL-based instruments, and replication in independent datasets.

Several next steps could strengthen mechanistic and translational conclusions. First, integrating tissue- and cell type-resolved data (single-cell transcriptomics, spatial profiling, and cell-state eQTLs) will clarify whether shared inflammatory signatures reflect true pathway activation within disease-relevant cells versus shifts in circulating immune composition. Second, incorporating orthogonal molecular layers such as proteomics (including cis-pQTL MR), methylation, and chromatin accessibility may help prioritize targets with consistent multi-omic support and improve causal inference. Third, expanding analyses to diverse ancestries and harmonized phenotypes will improve robustness and translational potential. Fourth, experimental validation using perturbation models (CRISPR perturb-seq, organoids, and in vivo aging models) should test whether modulating candidate targets (e.g., IL6R, TLR4, NLRP3, NOS2, GLP1R) recapitulates predicted pathway shifts and improves aging-relevant functional endpoints. Finally, clinical translation will benefit from developing standardized aging endpoints and biomarker strategies, including genetically informed patient stratification and target engagement metrics, to bridge the gap between genetic causality, molecular signatures, and therapeutic efficacy.

In summary, our integrative framework identifies conserved inflammatory and immune mechanisms linking aging and diverse ARDs, nominates shared and aging-specific targets with pathway-level coherence, and provides genetic support for a subset of candidates with potential for indication expansion. These findings support aging-related and pan-disease therapeutic strategies and motivate future work to refine target selection using tissue-resolved multi-omics, genetics-guided stratification, and experimental validation.

## Acknowledgements

We thank Ms. Elizaveta Ekimova for her technical assistance with figure design.

## References

1. Bischof, E., et al., Advanced pathological ageing should be represented in the ICD. Lancet Healthy Longev, 2022. 3(1): p. e12.

2. Organization, W.H. ICD-11 for Mortality and Morbidity Statistics. 2025 2025-01; Available from: https://icd.who.int/browse/2025-01/mms/.

3. McLaughlin, J., et al., OLS4: a new Ontology Lookup Service for a growing interdisciplinary knowledge ecosystem. Bioinformatics, 2025. 41(5).

4. Vamathevan, J., et al., Applications of machine learning in drug discovery and development. Nat Rev Drug Discov, 2019. 18(6): p. 463–477.

5. Eraslan, G., et al., Deep learning: new computational modelling techniques for genomics. Nat Rev Genet, 2019. 20(7): p. 389–403.

6. Zhavoronkov, A., et al., Deep learning enables rapid identification of potent DDR1 kinase inhibitors. Nat Biotechnol, 2019. 37(9): p. 1038–1040.

7. Pun, F.W., et al., Hallmarks of aging-based dual-purpose disease and age-associated targets predicted using PandaOmics AI-powered discovery engine. Aging (Albany NY), 2022. 14(6): p. 2475–2506.

8. Durinck, S., et al., Mapping identifiers for the integration of genomic datasets with the R/Bioconductor package biomaRt. Nat Protoc, 2009. 4(8): p. 1184–91.

9. Cerezo, M., et al., The NHGRI-EBI GWAS Catalog: standards for reusability, sustainability and diversity. Nucleic Acids Res, 2025. 53(D1): p. D998–D1005.

10. Deelen, J., et al., A meta-analysis of genome-wide association studies identifies multiple longevity genes. Nat Commun, 2019. 10(1): p. 3669.

11. Atkins, J.L., et al., A genome-wide association study of the frailty index highlights brain pathways in ageing. Aging Cell, 2021. 20(9): p. e13459.

12. Timmers, P.R., et al., Genomics of 1 million parent lifespans implicates novel pathways and common diseases and distinguishes survival chances. Elife, 2019. 8.

13. Vosa, U., et al., Large-scale cis- and trans-eQTL analyses identify thousands of genetic loci and polygenic scores that regulate blood gene expression. Nat Genet, 2021. 53(9): p. 1300–1310.

14. Perez, G., et al., The UCSC Genome Browser database: 2025 update. Nucleic Acids Res, 2025. 53(D1): p. D1243–D1249.

15. Hoffman, G.E. and E.E. Schadt, variancePartition: interpreting drivers of variation in complex gene expression studies. BMC Bioinformatics, 2016. 17(1): p. 483.

16. Leung, H., et al., Advancing Target Discovery Through Disease-Specific Integration of Multi-Modal Target Identification Models and Comprehensive Target Benchmarking System. bioRxiv, 2025.

17. Kamya, P., et al., PandaOmics: An AI-Driven Platform for Therapeutic Target and Biomarker Discovery. J Chem Inf Model, 2024. 64(10): p. 3961–3969.

18. de Magalhaes, J.P. and O. Toussaint, GenAge: a genomic and proteomic network map of human ageing. FEBS Lett, 2004. 571(1-3): p. 243–7.

19. Dolgalev, I., msigdbr: MSigDB Gene Sets for Multiple Organisms in a Tidy Data Format. 2025.

20. Korotkevich, G., et al., Fast gene set enrichment analysis. bioRxiv, 2021.

21. Yu, G. and Q.Y. He, ReactomePA: an R/Bioconductor package for reactome pathway analysis and visualization. Mol Biosyst, 2016. 12(2): p. 477–9.

22. Kolde, R., et al., Robust rank aggregation for gene list integration and meta-analysis. Bioinformatics, 2012. 28(4): p. 573–80.

23. Pun, F.W., et al., A comprehensive AI-driven analysis of large-scale omic datasets reveals novel dual-purpose targets for the treatment of cancer and aging. Aging Cell, 2023. 22(12): p. e14017.

24. Chang, C.C., et al., Second-generation PLINK: rising to the challenge of larger and richer datasets. Gigascience, 2015. 4: p. 7.

25. Hemani, G., et al., The MR-Base platform supports systematic causal inference across the human phenome. Elife, 2018. 7.

26. Giambartolomei, C., et al., Bayesian test for colocalisation between pairs of genetic association studies using summary statistics. PLoS Genet, 2014. 10(5): p. e1004383.

27. Ferrucci, L., et al., Serum IL-6 level and the development of disability in older persons. J Am Geriatr Soc, 1999. 47(6): p. 639–46.

28. Franceschi, C., et al., Inflamm-aging. An evolutionary perspective on immunosenescence. Ann N Y Acad Sci, 2000. 908: p. 244–54.

29. Harris, T.B., et al., Associations of elevated interleukin-6 and C-reactive protein levels with mortality in the elderly. Am J Med, 1999. 106(5): p. 506–12.

30. Ferrucci, L. and E. Fabbri, Inflammageing: chronic inflammation in ageing, cardiovascular disease, and frailty. Nat Rev Cardiol, 2018. 15(9): p. 505–522.

31. Liang, R., et al., The role of NLRP3 inflammasome in aging and age-related diseases. Immun Ageing, 2024. 21(1): p. 14.

32. Li, X., et al., Inflammation and aging: signaling pathways and intervention therapies. Signal Transduct Target Ther, 2023. 8(1): p. 239.

33. Yamamoto, R., et al., Tissue-specific impacts of aging and genetics on gene expression patterns in humans. Nat Commun, 2022. 13(1): p. 5803.

34. Cao, W., IFN-Aging: Coupling Aging With Interferon Response. Front Aging, 2022. 3: p. 870489.

35. Calabrese, V., et al., Aging and Parkinson’s Disease: Inflammaging, neuroinflammation and biological remodeling as key factors in pathogenesis. Free Radic Biol Med, 2018. 115: p. 80–91.

36. Loeser, R.F., Aging processes and the development of osteoarthritis. Curr Opin Rheumatol, 2013. 25(1): p. 108–13.

37. Stegeman, R. and V.M. Weake, Transcriptional Signatures of Aging. J Mol Biol, 2017. 429(16): p. 2427–2437.

38. Gan, C., et al., Liver diseases: epidemiology, causes, trends and predictions. Signal Transduct Target Ther, 2025. 10(1): p. 33.

39. Interleukin-6 Receptor Mendelian Randomisation Analysis, C., et al., The interleukin-6 receptor as a target for prevention of coronary heart disease: a mendelian randomisation analysis. Lancet, 2012. 379(9822): p. 1214–24.

40. Ferreira, R.C., et al., Functional IL6R 358Ala allele impairs classical IL-6 receptor signaling and influences risk of diverse inflammatory diseases. PLoS Genet, 2013. 9(4): p. e1003444.

41. Bonafe, M., et al., A gender--dependent genetic predisposition to produce high levels of IL-6 is detrimental for longevity. Eur J Immunol, 2001. 31(8): p. 2357–61.

42. Rosa, M., et al., A Mendelian randomization study of IL6 signaling in cardiovascular diseases, immune-related disorders and longevity. NPJ Genom Med, 2019. 4: p. 23.

43. Wissel, S., et al., Genetic interleukin-6 receptor blockade, Chronic Disease Risk and Longevity. Results from the Women’s Health Initiative. Eur J Prev Cardiol, 2025.

44. Singh, D., et al., Selective inducible nitric oxide synthase inhibition has no effect on allergen challenge in asthma. Am J Respir Crit Care Med, 2007. 176(10): p. 988–93.

45. Hoivik, H.O., et al., Lack of efficacy of the selective iNOS inhibitor GW274150 in prophylaxis of migraine headache. Cephalalgia, 2010. 30(12): p. 1458–67.

46. Papapetropoulos, A., A.J. Hobbs, and S. Topouzis, Extending the translational potential of targeting NO/cGMP-regulated pathways in the CVS. Br J Pharmacol, 2015. 172(6): p. 1397–414.

47. Li, C.Y., et al., Repurposing nitric oxide donating drugs in cancer therapy through immune modulation. J Exp Clin Cancer Res, 2023. 42(1): p. 22.

48. Burnett, A.L., The role of nitric oxide in erectile dysfunction: implications for medical therapy. J Clin Hypertens (Greenwich), 2006. 8(12 Suppl 4): p. 53–62.

49. Tewari, D., et al., Role of Nitric Oxide in Neurodegeneration: Function, Regulation, and Inhibition. Curr Neuropharmacol, 2021. 19(2): p. 114–126.

