## Supplementary Figures for "Integrating AI and causal genetics to prioritize therapeutic targets for aging and age-related diseases"

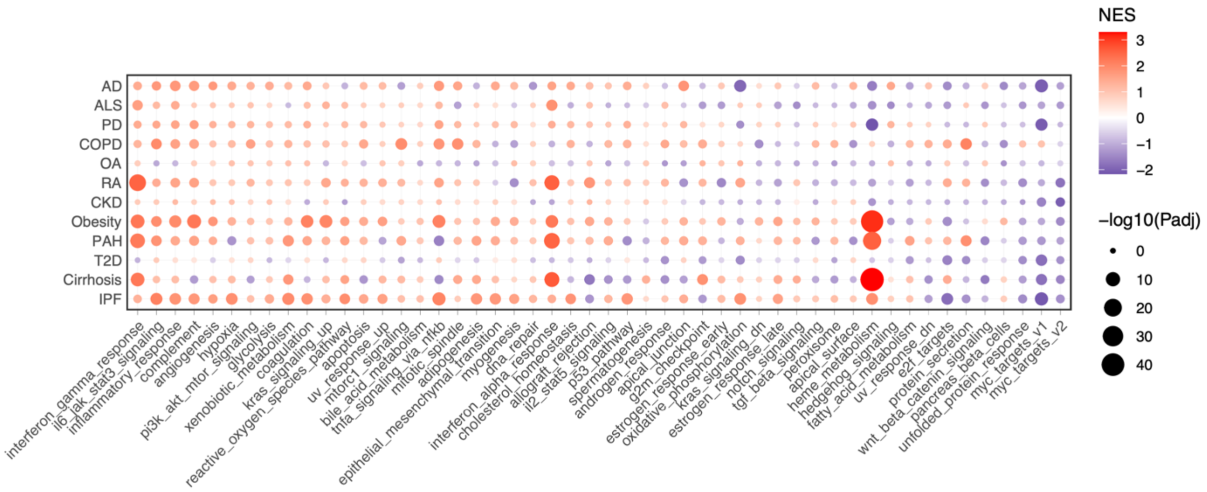


Supplementary Figure 1. Perturbation of the 50 analyzed hallmark gene sets in 12 ARDs. Red, NES > 0; Blue, NES < 0. NES, Normalized enrichment score.

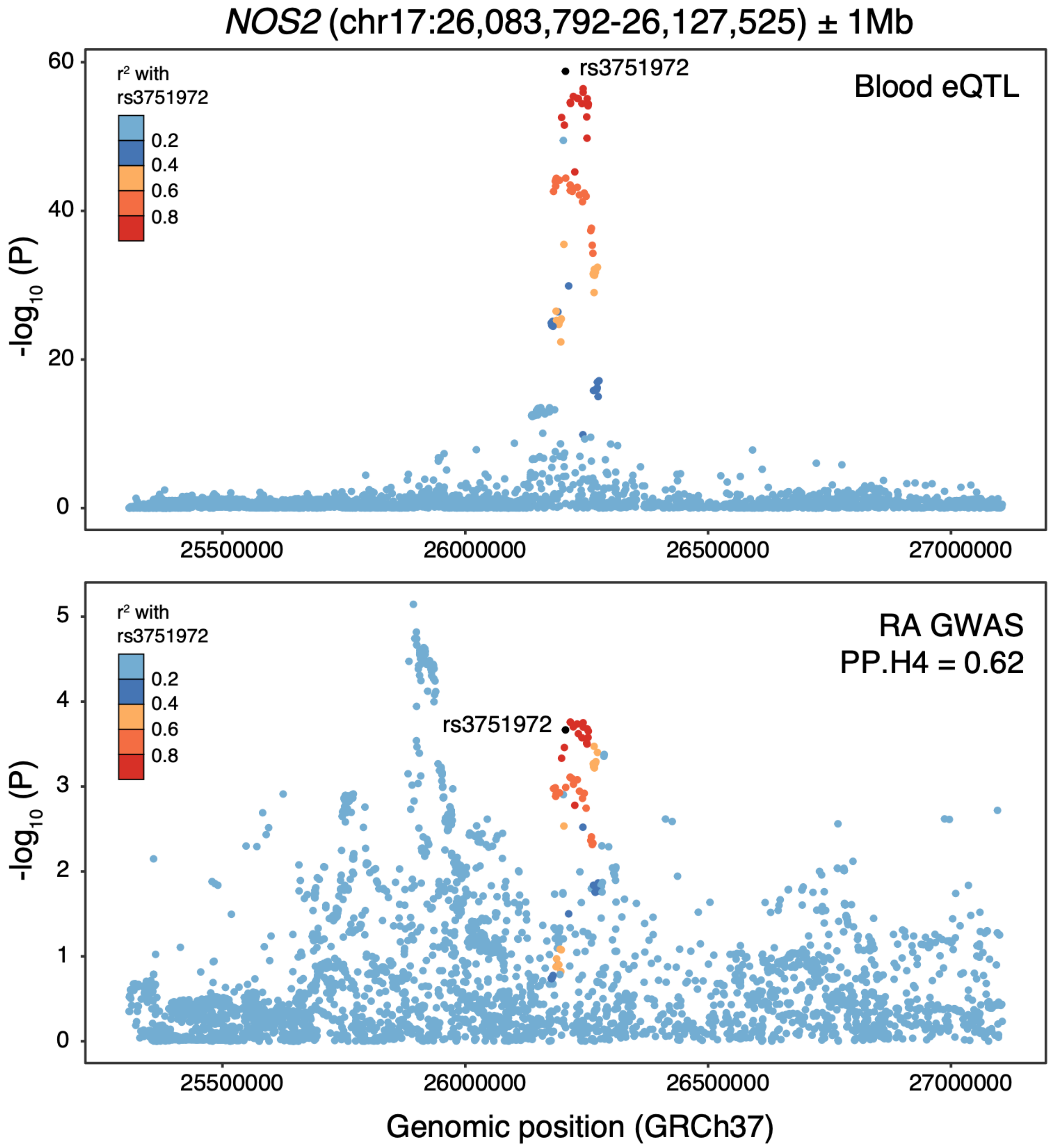


Supplementary Figure 2. Colocalization analysis showing moderately shared genetic signals between *NOS2* expression and rheumatoid arthritis at the *NOS2* locus.
